# Novel Dual Labeled Fluorescence Probe Based Assay to Measure the Telomere Length

**DOI:** 10.1101/507616

**Authors:** Itty Sethi, Gh. Rasool Bhat, Rakesh Kumar, Ekta Rai, Swarkar Sharma

## Abstract

Telomeres are highly repetitive regions capping the chromosomes and composed of multiple units of hexa-nucleotides, TTAGGG, making their quantification difficult. Most of the methods developed to estimate telomeres are extensively cumbersome and expensive. The quantitative polymerase chain reaction (qPCR) based assay is relatively easy and cheaper method that applies SyBr Green dye chemistry to measure telomere length. As SyBr Green dye fluoresces after intercalation into the dsDNA, lack of differentiation between specific PCR target products and unspecific products is a limitation and it affects accuracy in quantitation of telomeres. To overcome the limitations of SyBr Green, we developed a dual labeled fluorescence probe based quantitative polymerase chain reaction (qPCR) to measure the telomere length. This robust, accurate and highly reproducible (R^2^=0.96) proprietary method (patent pending), yet cost effective and easy, utilizes a probe that targets specifically the telomeric DNA.

## Introduction

Telomeres are the prominent structures capping the ends of chromosomes. These are composed of tandom hexa-nucleotide repetitive sequences, TTAGGG, in humans and terminate at 3′ single strand guanine overhang^1^. The environmental impact on telomere length leads to several chronic, stress induced, metabolic and aging disorders^2^. In modern era, telomere length has become a biomarker of cell senescence^3^. Subsequently, the interest of measuring the telomere length has increased in biomedical, clinical, and commercial field. This leads the investigators to explore various methods to measure telomere length. The conventionally well appreciated method is telomere length restriction fragment length (TRF) analysis^4^. Besides this, other methods include Single Telomere Length Analysis (STELA), Fluorescent in situ Hybridization (FISH), Quantitative PCR (qPCR), Monochrome Multiplex Quantitative PCR (MMQPCR), Absolute Quantitative PCR (aqPCR), and sequencing^5, 6^. All the techniques have their own advantages and limitations. However, qPCR is proposed to be a better version and best suited to large epidemiologic studies, as it requires less DNA (5-20ng) and is relatively less time consuming.

MMQPCR method is based on amplification of telomeric region by a unique set of telomere primers in such a way that a fixed length of telomere product is obtained which is then compared to the single copy gene (scg) product^7^ (Figure 1). Though MMQPCR is a multiplex method yet amplification and detection of both products (telomere and single copy gene) may not be possible at the same time point because of the assay design (details in methodology). The other limitation is the use of SyBr Green dye chemistry as detection system. The disadvantage of SyBr Green based assays is it’s binding to any ds DNA which is not target specific. Owing to its unspecific nature of binding, there are chances of SyBr outflow resulting in inconsistencies. The copy number difference should be relatively very high between the single copy gene and telomeres based on the fact that normal healthy human genome (excluding aneuploidies and various other pathologies) has at least 46 copies of chromosomes and telomeres cap the ends of all the chromosomes. However, we did not find it by SyBr Green based MMQPCR method to estimate telomeres. We experienced that SyBr Green outflows from telomere region to single copy gene readings in this method. Thus, we concluded that the adoption of the present MMQPCR method is also very challenging.

**Figure 1:**
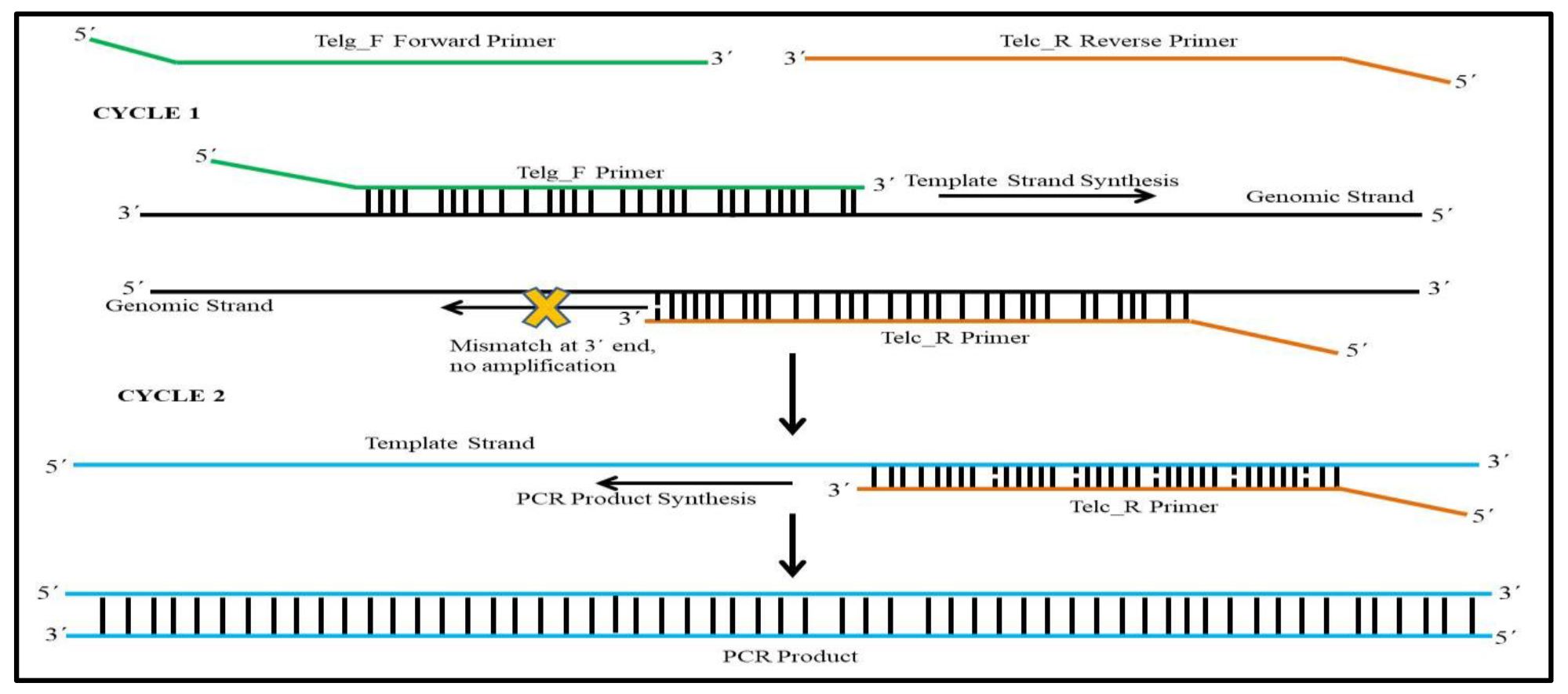
Schematic representation of working mechanism of telomere primers. In cycle Telg_F (forward primer) binds to the telomeric DNA and amplifies the template strand. Telc_R (reverse primer) binds to the telomeric DNA but cannot amplify the sequence due to a specific mismatch at the 3′ end. In cycle 2, Telc_R (reverse primer) binds to the template strand and produces the final telomere product of fixed length, 79bp.

The main objective of this work was to establish a method that exploits the versatility and easiness of qPCR to telomere measurement, for large sample size studies especially, but at the same time with highest level of specificity, robustness and replicability. We believe we have achieved this by developing a novel dual labeled fluorescent probes-based assay [also called as TaqMan assay^8^ and universally known for their specificity^9^] and method for telomere length measurement in an innovative way. This method allows the accurate capturing of the fluorescence signal from the target regions by including additional proprietary components and steps (detailed and discussed subsequently) in MMQPCR methodology^7^.

## Methods

The methodology involved use of human DNA samples thus, all experiment procedures were conducted according to the guidelines and regulations set by Institutional Ethics Review Board (IERB), Shri Mata Vaishno Devi University (SMVDU). The use of anonymous and unidentified human DNA samples for this work has also been approved by the IERB, SMVDU. All the samples used in the present study had been collected with a written informed consent to provide 2ml of blood by venipuncture. A conventional phenol-chloroform method and FlexiGene^®^ DNA kit, QIAGEN (catalogue No. 51206) method was used to extract genomic DNA from the blood samples. The quantity and quality of the genomic DNA was analyzed by performing UV spectrophotometer (eppendorf Biospectrometer®, Hamburg Germany) analysis and Gel electrophoresis. The genomic DNA was diluted to a concentration of 10ng/μl and stored at 4°C till further use. Leukocyte Telomere Length was assessed by using Monochrome Multiplex Quantitative Polymerase Chain Reaction (MMQPCR) method^7^ as well as by our dual labeled fluorescence probe based assay. All samples were run in triplicate to check the relative telomere length and access the reproducibility of the results.

### MMQPCR Methodology

To begin with, all the samples were run in triplicate with reported methodology^7^ for MMQPCR but findings were inconsistent as described in results. Thus, we tried different PCR cycling conditions and carried out experiments. The final modified conditions that worked best are provided. Stage 1: single cycle of 95°C for 15 minute; Stage 2: 2 cycles of 94°C for 15 seconds, 49°C for 1 minute; Stage 3: 5 cycles of 85°C for 20 seconds, 59°C for 30 seconds; Stage 4: 30 cycles of 94°C for 15 seconds, 59°C for 30 seconds with the signal acquisition for telomere product, 84°C for 30 seconds and 85°C for 20 seconds with the signal acquisition for single copy gene product; Stage 5: a single default cycle (Agilent Mx3005p) of dissociation curve - 95°C for minute, 55°C for 30 seconds and 95°C for 30 seconds. A total of 50 healthy individual samples were run in triplicates. The final concentration of reagents in each of 20 μl MMQPCR reaction mixture were 1X SYBR Green Master mix (Applied Biosciences), 0.6μM telg_F primer, 0.3μM telc_R primer, 0.5 μM albu_F primer, 0.5 μM albd_R primer, 2% DMSO, 0.8M Betaine (Invitrogen) and ~20ng DNA. The PCR efficiency was analyzed by executing a standard curve of 3 fold serial dilution, spanning 81-fold range (151ng, 51.7ng, 16.1ng, 6.1ng and 1.7ng DNA). The standard curve experiment was performed separately for both telomere and single copy gene in a same reaction plate independently (single plex) (Supplementary Figure 5-10).

### Dual Labeled Fluorescence Probe Based Assay Design and Conditions

A novel specific dual labeled fluorescence probes based assay was developed. Probes were designed to detect the telomere product obtained by unique telomere primer set as described in figure 1^7^. This assay overcomes the limitations of SyBr Green and increases the robustness and reproducibility of the telomere measurement by qPCR. Two set of probes and methods were designed. The probe consisted of reporter at 5′ and quencher at 3′ (Figure 2 & 7). The first set has telomere probe (Tel_P_a_) of size 21 bases and FAM as reporter dye and BHQ as quencher whereas WISP3 as single copy gene with probe WISP3_P of size 24 bases in exon 5 of WISP3 gene. In this single copy gene probe, HEX was used as a reporter dye and BHQ as quencher. The second set was developed to improve the assay. It was composed of 27 bases probe, Tel_P_b_ with FAM as reporter dye and BHQ as quencher. This time, albumin was used as single copy gene and probe ALB_P of size 29 bases was designed. HEX was used as a reporter dye and BHQ as quencher for this probe, ALB_P (Figure 2–4). Primers and probes were synthesized commercially by IDT.

**Figure 2:**
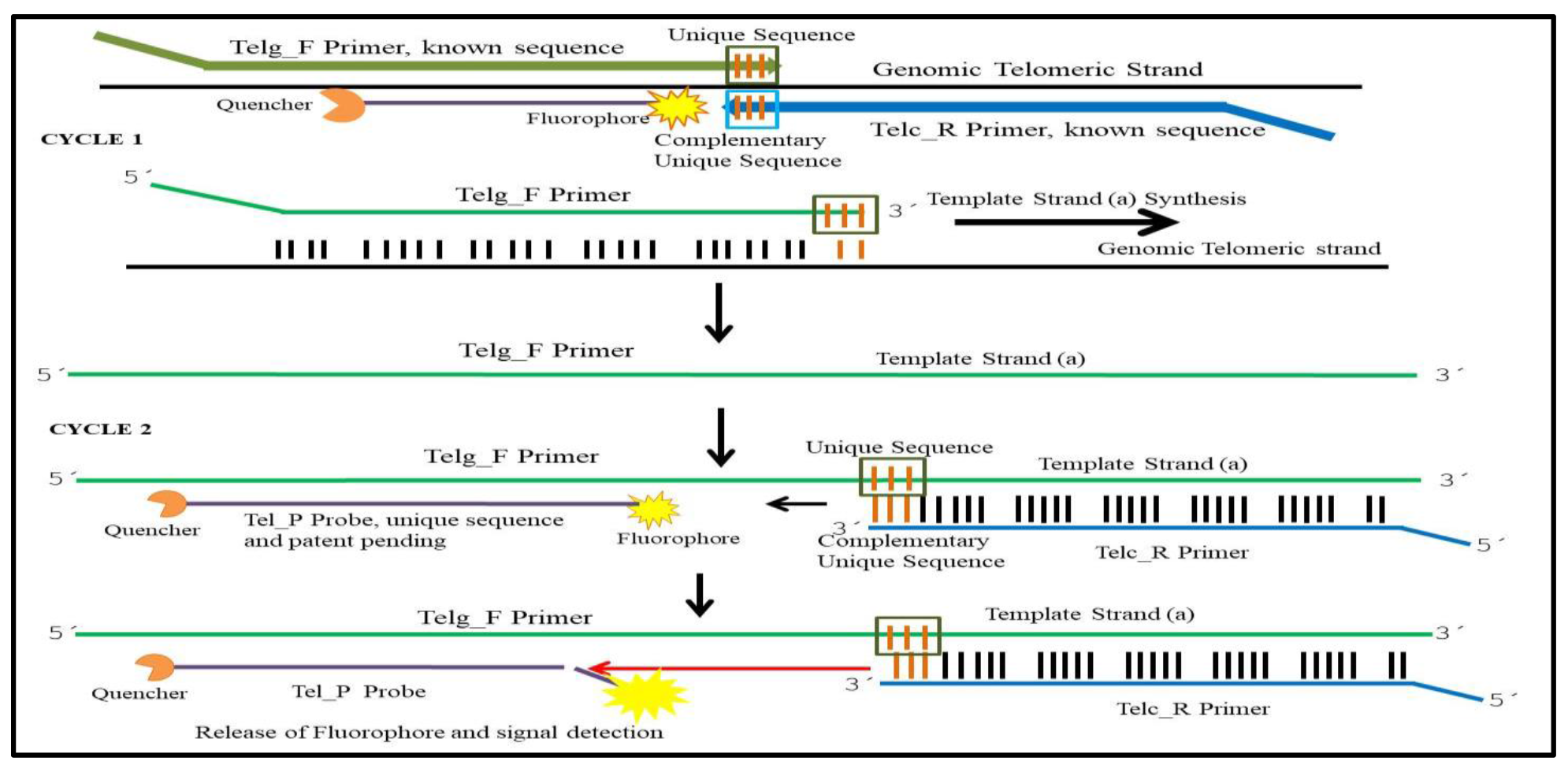
Schematic representation of dual labeled fluorescence probe based assay (using Tel_P probe) for Telomere product. Cycle 1 represents the first step of PCR where the Telg_F forward primer binds to the telomeric DNA and synthesis of template (a) for telomere assay takes place; Cycle 2 represents the second step in which Telc_R reverse primer and Tel_P probe anneals to this template strand (a), the product of cycle 1. Fluorescence is produced after the fluorophore gets cleaved off by polymerase while forming the final product. The brown colored lines (nucleotides) represent the unique sequence, located on Telg_F forward and Telc_R reverse primer which are complementary to each other. These are the bases where Telc_R reverse primer binds with the template strand (a) and precedes the reaction for final product formation and signal production.

**Figure 3:**
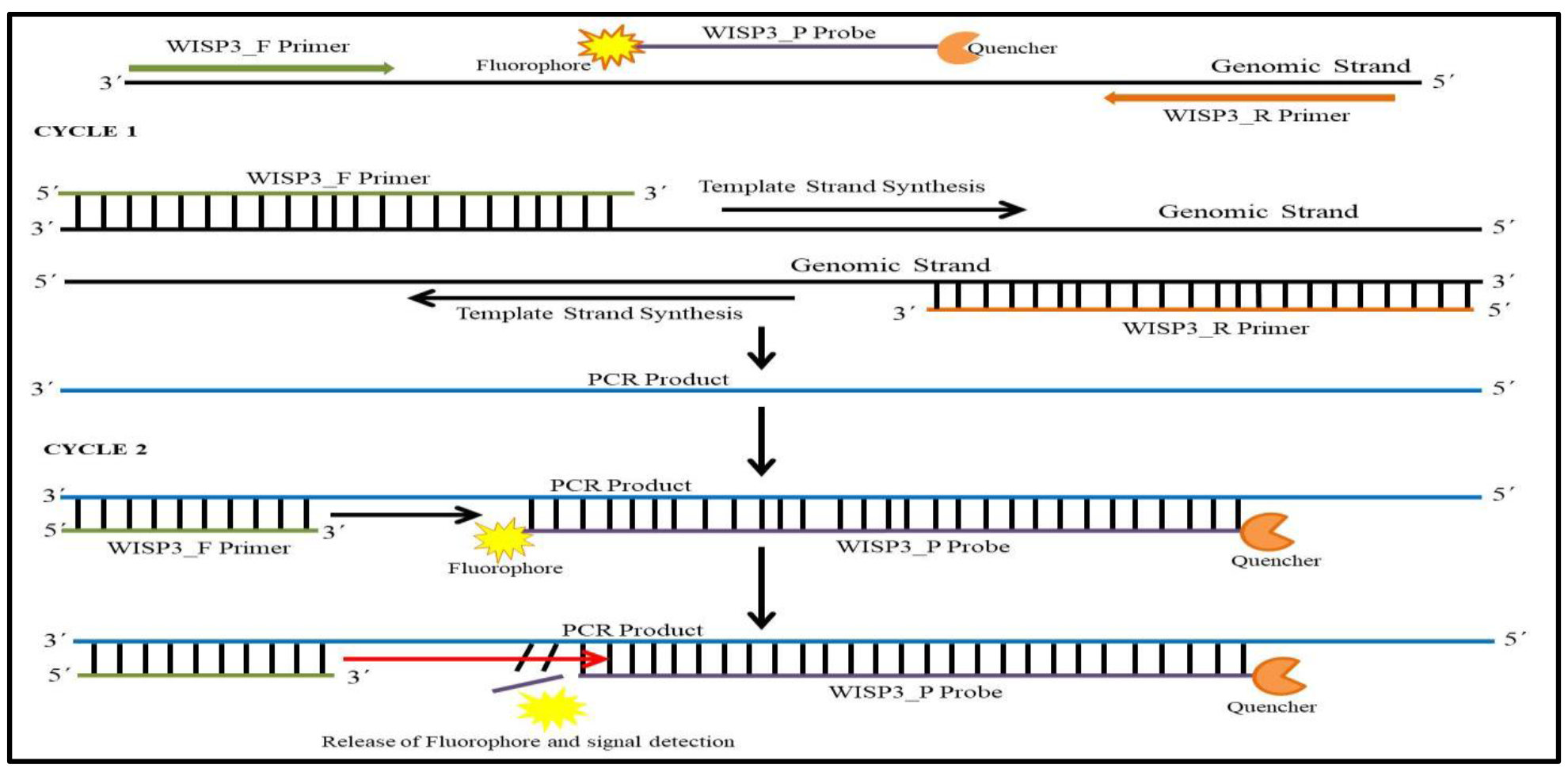
Schematic representation of innovative dual labeled fluorescence probe based assay for WISP3 product. Cycle 1 represents the first step of PCR where the WISP3_F forward primer and WISP3_R reverse primer anneals to the target sequence and forms the desired product which in second cycle (Cycle 2) later used as target sequence where the WISP3_P probe anneals and cleaved off by polymerase thereby giving the florescence.

**Figure 4:**
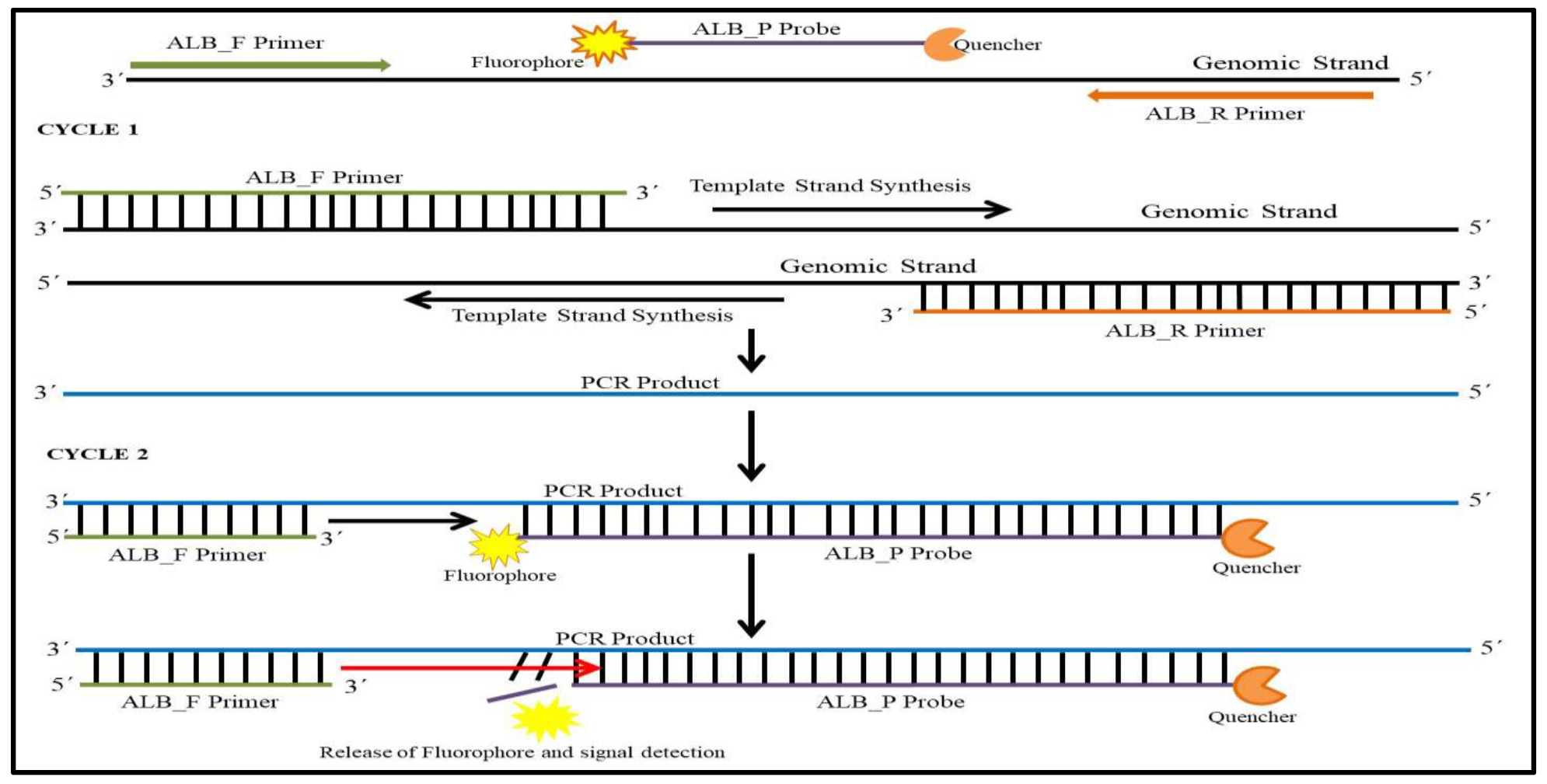
Schematic representation of dual labeled fluorescence probe based assay for single copy gene (Albumin) product. Cycle 1 represents the first step of PCR where the ALB_F forward primer and ALB_R reverse primer anneals to the target sequence and forms the desired product which in second cycle (Cycle 2) later used as target sequence where the ALB_P probe anneals and cleaved off by polymerase thereby giving the florescence.

**Figure 5:**
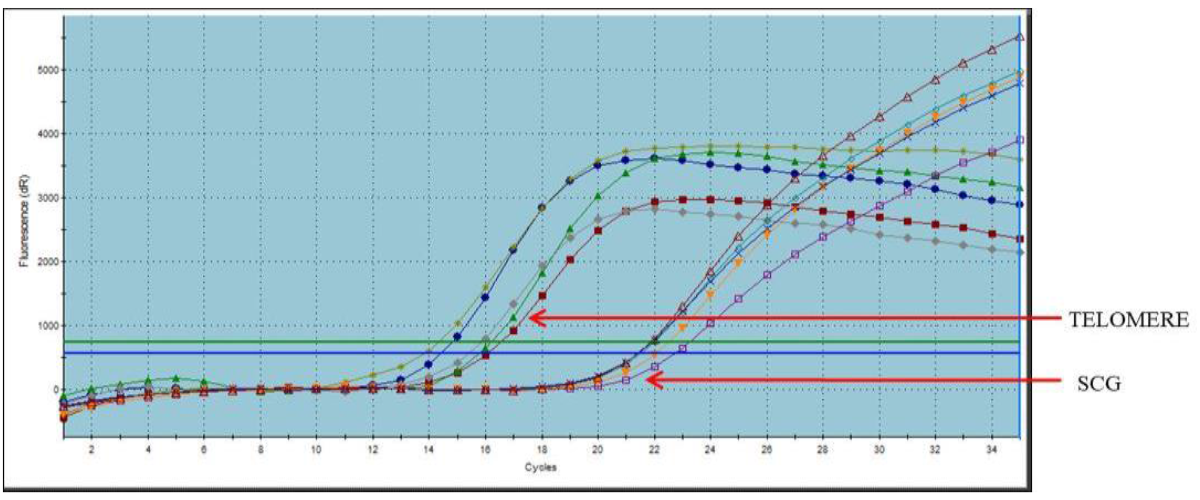
Amplification plot of telomere and albumin product generated from dual labeled fluorescence probe based assay. Telomere product crosses threshold at 14-16 cycles and single copy gene (albumin) product crosses threshold at 22-23 cycles. The difference of Ct value between the telomere and single copy gene is about 8-9 cycles.

### Conditions used with first set of probes (Tel_P_a_ and WISP3_P)

The final concentration of reagents in 10 μl qPCR reaction mixture were 1X PCR Buffer (Sigma) with 1.5-2.0mM MgCl_2_, 0.4μM Telg_F primer, 0.2μM Telc_R primer, 0.3μM Tel_P probe, 0.5 μM WISP3_F primer, 0.5 μM WISP3_R primer, 0.5μM WISP3_P probe, 0.4mM dNTPs (Sigma), 0.5U Taq Polymerase (Sigma) and 10ng DNA. Agilent Mx3005p Stratagene Real Time was used to perform the assay. All the samples were evaluated in triplicates. We performed the assay on a total of 25 healthy individual samples, overlapping with MMQPCR to compare the results. Of these 23 samples worked well and were further used for the analyses. Multiple PCR conditions were used to standardize and finalized quantitative Real Time PCR conditions were: Stage 1: single cycle of 95°C for 4 minute; Stage 2: 6 cycles of 95°C for 15 seconds, 52°C for 15 seconds, and 72°C for 30 seconds; Stage 3: 2 cycles of 95°C for 4 minutes, 95°C for 15 seconds, and 49°C for 1 minute; 38 cycles of 90°C for 30 seconds, 52°C for 15 seconds, and 54°C for 1 minute, with the signal acquisition at 54°C for both the probes (telomere and single copy gene).

### Conditions used with second set of probes (Tel_P_b_ and ALB_P)

The final concentration of reagents in 10 μl qPCR reaction mixture were 1X PCR Buffer (Sigma) 1.5-2.0mM MgCl_2_, 0.4μM Telg_F primer, 0.2μM Telc_R primer, 0.3μM Tel_P probe, 0.5 μM ALB_F primer, 0.5 μM ALB_R primer, 0.5μM ALB_P probe, 0.4mM dNTPs (Sigma), 0.5U Taq Polymerase (Sigma) and 10ng DNA. Agilent Mx3005p Stratagene Real Time was used to perform the assay. All the samples were evaluated in triplicates and we performed the assay on a total of 210 samples in two independent sets of size 120 and 90. Quantitative Real Time PCR conditions were finalized after testing various thermal cycling conditions. Stage 1: single cycle of 95°C for 5 minute; Stage 2: 5 cycles of 95°C for 15 seconds, 49°C for 1 minute; Stage 3: 35 cycles of 85°C for 30 seconds, 49°C for 30 seconds, and 59°C for 1 minute, with the signal acquisition at 59°C for both the probes (telomere and single copy gene).

The PCR efficiency of the dual labeled fluorescence probe based assay was analyzed by performing a standard curve of 3 fold serial dilution, spanning 81-fold range (190ng, 63.33ng, 21.11ng, 7.04ng and 2.35ng DNA). The standard curve experiment was performed separately for both telomere and single copy gene in a same reaction plate, independently (single plex) (Figure 6 and Supplementary Figure 13-14).

**Figure 6:**
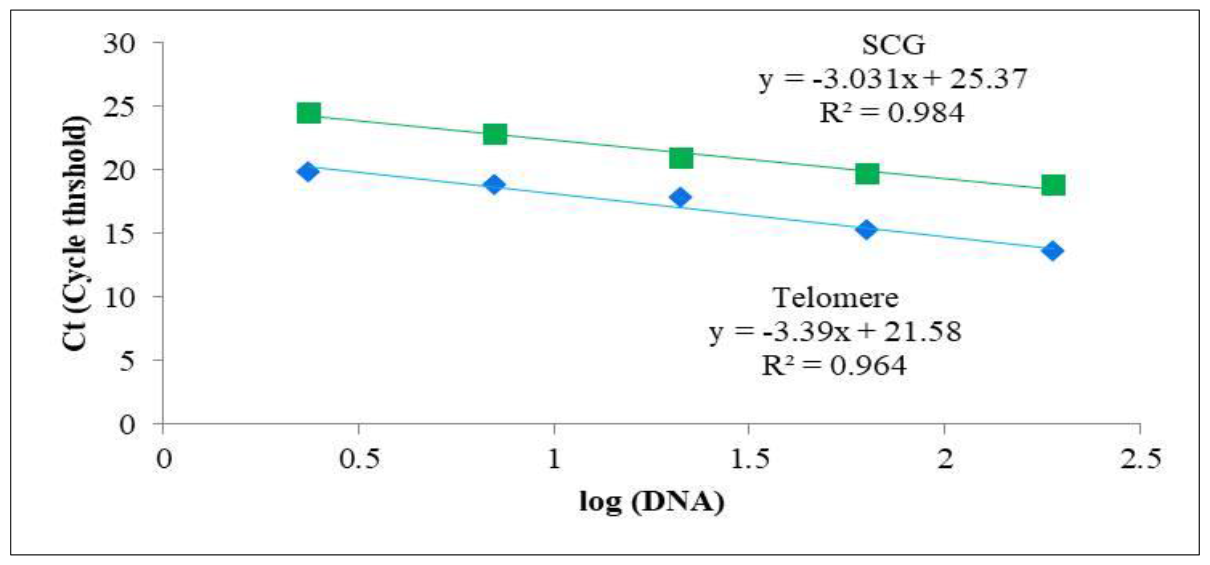
Schematic representation of standard curve for telomere and scg product step 3 (59°C) of stage 3 with R^2^= 0.96 and 0.98, slope= -3.39 & -3.03 and efficiency of 97.2% & 113.8% respectively, generated by dual labeled fluorescence probe based assay methodology.

### Data Collection and Analyses

The data was exported in the excel format from the MxPro Software. The triplicates were averaged and then the averaged Ct value was used for further calculation of relative telomere length as T/S ratio. T/S ratio was calculated on the basis of Ct values, as [2Ct (telomeres)/2Ct (scg)]^-1^ = 2^-ΔCt^ ^10^. T/S ratio is relative estimation of telomere copies as compared to the single copy gene in an individual sample that corresponds to the relative telomere length.

### Gel Electrophoresis

PCR product was analyzed by performing 3% agarose gel electrophoresis and 10% native Poly acrylamide gel electrophoresis (PAGE), whenever required.

## Results and Discussion

The working of telomere primers in MMQPCR method is different than the routine primer methodology. In our experiment, the telg_F (forward primer) first anneals to its target sequence (telomeric DNA) and produces the first product of qPCR. Telc_R (reverse primer) is complimentary to the Telg_F primer product and anneals to this sequence giving the final product of 79bp (Figure 1). According to established MMQPCR protocol^7^, two amplified products were expected; the fixed length telomere product (79bp) and single copy gene (albumin) product of 98bp. However, we observed that the dissociation curve peak was varying (very low) for telomere product as well as visualization of two amplified products in gel was also inconsistent (Supplementary Figures 1–5). Even the difference between the Ct (threshold) values of two amplified products, telomere and single copy gene, was less. These low Ct value differences imply very less difference of copy numbers between telomere and single copy gene. Despite the fact that telomeres exist in multiple folds compared to the single copy gene, it is not statistically accurate (Supplementary Table 1).

We modified the thermal cycling profile described as in the methodology section and validated our conditions with the gel electrophoresis (Supplementary Figure 1). A 3-fold serial dilution linear curve of log DNA concentration versus the Ct (cycle threshold) values was also produced for products, telomere and single copy gene (Supplementary Figure 5-10). The observations for the difference in Ct values of the two products improved only a bit (Supplementary Figure 2-3), and the telomere product dissociation curve peak was still inconsistent (Supplementary Figure 4). We hypothesized the difference in Ct values was less because of the leakage of SyBr Green signal from telomere to single copy gene but were not able to comprehend it. To understand the reason better, we explored extensively about the expected product formed at telomere cycle. Interestingly, we noted that though the telomere amplification and signal is acquired at 59°C, the melting temperature of amplified 79bp product is 87.4 (tm adjusted for salt presence expected in PCR buffer reagents^11^). We got justification for our hypothesis that existence of this 79bp fragment after couple of cycles, provides for SyBr Green to bind and produce signal at single copy gene estimation time point. This thereby increases the fluorescence signal of single copy gene and reduces the Ct value difference (Supplementary Figure 2-4).

With this background, a better telomere region targeting method was designed. We present here a novel probes-based assay that overcomes the limitations of previous methodologies and uses unique probes that were specifically developed to target telomere region. Thus, provides a better method to analyze the telomere length efficiently and accurately. The method is based on the principle of dual labeled fluorescent probes based assays, which consists of a fluorescent reporter and quencher attached to the 5’ end and 3’ end of the probe, respectively^9^. It further relies on the 5′-3′ exonuclease activity of DNA polymerase^12^. Taq polymerase while extending the primer, releases the reporter attached to the 5′ end of the probe thereby separating it from the quencher during the PCR and produces the desired signal when quencher is no longer able to quench the fluorescence of reporter (Figure 2 and 7). In this assay, both the products (telomere and single copy gene) are easily identifiable irrespective of each other’s amplification because of unique probes and fluorescent dyes. Single copy gene primers and probe are designed according to the set standards. Primers to target telomere region were adapted^7^ (Figure 1) and unique telomere probes, Tel_P_a_ and Tel_P_b_ probes were designed to work specifically in the reaction. The unique telomere probes Tel_P_a_ (21 bases) and Tel_P_b_ (27 bases) were designed in a way to anneal exactly to the Telg_F forward primer product (Figure 2). These probes were designed in a fashion that last appropriate number of bases (proprietary information) are free, to ensure the binding of polymerase, specific extension and excision of the reporter from the probe to produce the desired signal for quantification. The methodology partially adopts steps as described^7^. Firstly the Telg_F primer anneals to the telomeric DNA and template gets synthesized. The Telc_R primer then anneals to the Telg_F primer product in a specific manner and at the same time probe anneals to its target sequence (product of Telg_F primer extension) (Figure 1 & 2). After annealing of both probe and primer, polymerase extends the reverse primer and also excises the reporter, by its 5′-3′exonuclease activity, thereby producing fluorescence. We validated the specific amplification by 3% agarose gel electrophoresis and observed fragments of anticipated sizes (Supplementary Figure 11). As anticipated, the difference in the Ct values between telomere and single copy gene in individual samples improved to an average of 6-9 cycles and observed to go up to 12 cycles (Figure 5 and supplementary figure 12). Further, 3-fold serial dilution linear curve of log DNA concentration versus the Ct (cycle threshold) values was obtained for both telomere and single copy gene to analyze the PCR efficiency and the curve were generated for both the products (Figure 6 & Supplementary Figure 13-14). The coefficient of determination (R^2^=0.96) and efficiency for the standard curve 97.2% implied highly replicative and accurate quantitation in all the samples.

**Figure 7:**
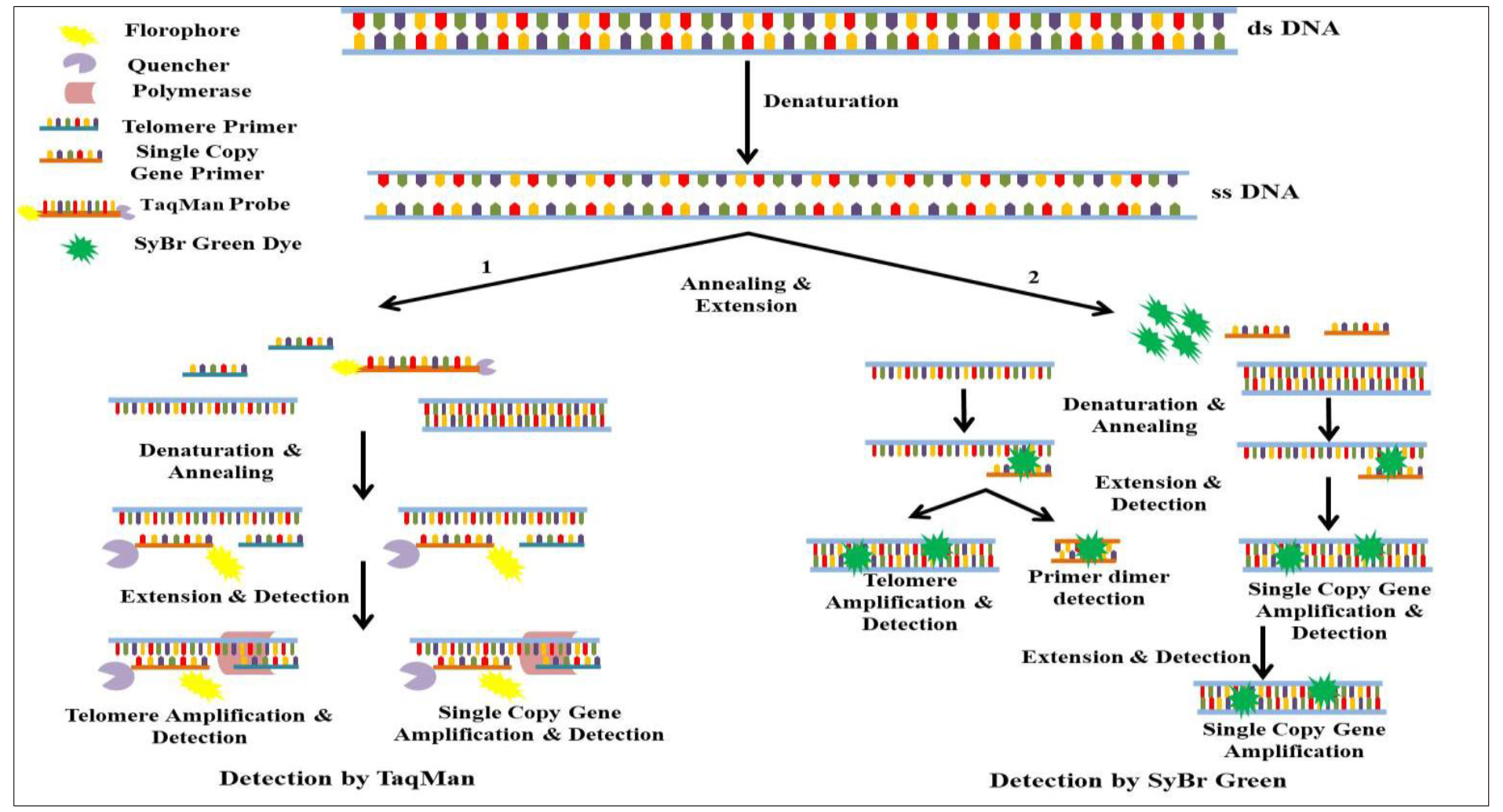
Schematic representation of difference between dual labeled fluorescence probes based assay & SyBr Green (MMQPCR). Panel 1 - describes the dual labeled fluorescence probe based assay method where the dual labeled fluorescence probe anneals to its specific target region and the fluorophore gets cleaved off by the polymerase resulting in the increase of fluorescence intensity with each cycle of PCR. Panel 2 - describes the SyBr Green dye mechanism where SyBr Green intercalates in the ds DNA. As it is an intercalating dye and the fluorescence intensity increases with each PCR cycle; it intercalates in the ds DNA without distinguishing between the specific and unspecific products.

We compared this method with MMQPCR on total of 23 samples and found noteworthy difference in the average delta Ct values obtained from both the methods. The delta Ct value is estimated as the difference between the Ct value of telomere and single copy gene (ΔCt = ΔCt_Telomere_ - ΔCt_Single copy gene_). In case of MMQPCR, the delta Ct value was -0.44 while in dual labeled fluorescence probe based assay, it was -5.04 in the same sample set. The negative value indicates higher numbers of cycles are required to amplify single copy gene as compared to telomere. We emphasize the substantial difference in the Ct values, observed in our assay as expected but not obtained by MMQPCR, is because of the precision and robustness of this dual labeled fluorescence probe-based assay (Supplementary Table 2).

Tel_P_b_ probe was designed to improve our assay further. Additionally, we used albumin as single copy gene this time and designed the primer set (ALB_F and ALB_R) and probe (ALB_P) in specific region and manner (proprietary information, Patent pending) to validate dual labeled fluorescence probe based telomere assay, simultaneously. The difference between the first Telomere probe (Tel_P_a_) and second Telomere probe (Tel_P_b_) is only of 6 bases, but it was introduced to make it more specific, accurate and robust. To test implications and robustness of Tel_P_b_ probe and albumin as single copy gene, we screened the two independent human DNA sample cohorts from population of Jammu and Kashmir. Cohort 1 had n=120 and cohort 2 had n = 90, human DNA samples. The average delta Ct value (indicates fold difference between telomere and single copy gene) in case of cohort 1 was observed to be 5.42 while in cohort 2, it was 5.84 (Supplementary Table 3 and Supplementary Figure 13). The copy number difference in both the independent sets further strengthened our observations. Therefore, we inferred from the results that our telomere probes are unique and specific.

## Conclusion

We conclude that this dual labeled fluorescence probe assay based methodology to estimate telomere length is better than SyBr Green based assays. We find explanation in the fact that intercalation of SyBr Green dye between the ds DNA without discrimination between the target DNA and unspecific products causes inconsistent results (Figure 7). Further, we believe that leakage of telomere based SyBr Green fluorescence at single copy gene estimation point is reducing the anticipated difference between telomere and single copy gene which is potentially a major limiting factor of the MMQPCR assay^7^. The designing of probes in telomere region is perplexing than the development of routine probes as telomeres are extensive repeat regions. It is potentially the main reason that no one has tried to develop the dual labeled fluorescence probes for telomeres. However, we exploited the existing knowledge^7^, for the purpose of amplifying the region specifically, and by introducing advancement and novelty to this approach rectified these limitations. Our dual labeled fluorescence probe based method allows the capture of fluorescence acquisition from the target region robustly and specifically. We anticipate that this method could be easily adapted by researchers, with flexibility to perform it on any Real Time PCR based platform. This will help extensively in telomere estimation based studies, especially where bulk screening of the samples is required.

## Supporting information

Supplementary Data_Sethi et.al

## Acknowledgment

SS acknowledges grant from DST-SERB, GoI (SB/YS/LS-91/2015). SS, ER and IS acknowledge the financial assistantship to IS from SMVDU for her Ph.D. thesis work. RK acknowledges grant from DST, GoI project RP-93 (DST/SSTP/J&K/459). RK and GRB acknowledge the financial assistantship to GRB from DST, GoI project RP-93 (DST/SSTP/J&K/459).

## Author Contributions

SS, IS and GRB conceived the concept, designed the work plan and prepared the MS. SS, ER and RK provided the samples for experimental work. IS and GRB performed the experiments and IS, GRB and SS carried out the data analyses. All the authors contributed in finalization of the MS.

## Competing Interests

The authors declare financial competing interests as patent related to work and content in MS is filed with Indian Patent office, GoI and granting of patent is pending.

## Supplementary Tables and Figure Legends

**Supplementary Table 1:** MMQPCR performed on 50 samples, the average delta Ct observed was -0.44. This implies the difference between copy number of telomere and single copy gene is very low.

**Supplementary Table 2:** Comparison of both methodologies - MMQPCR and dual labeled fluorescence probe based assay for Telomere length measurement in a same sample set of 25. The samples were run in triplicates. Two of the samples did not provide the triplicate results hence removed from analysis. The copy number difference estimated by dual labeled fluorescence probe based assay between the telomere and single copy gene was 5.04 folds as compared to 0.44 estimated by MMQPCR assay.

**Supplementary Table 3:** Telomere length Measurement by dual labeled fluorescence probe based assay in two independent cohorts of size n=120 and n=90 respectively, from population of Jammu and Kashmir, India. Samples were run in triplicates and averaged Ct values were used for each sample to estimate average ΔCt values (ΔCt = Ct_Telomere_ - Ct_Single copy gene_).

**Supplementary Figure 1:** 10% Native PAGE was performed to analyze the product obtained from MMQPCR; Telomere product was of 79bp and single copy gene product was of 98bp. L1 represents 50bp ladder, L2 represents albumin and telomere product, L3 represents telomere product, L4 represents albumin product, L5 represents albumin and telomere product, L6 represents telomere product, L7 represents albumin product and L8 represents NTC.

**Supplementary Figure 2:** Representation of amplification plot of telomere product obtained from MMQPCR. Telomere product crossed the threshold after 19 cycles of PCR.

**Supplementary Figure 3:** Representation of amplification plot of Single Copy Gene (albumin) product obtained from MMQPCR. Single copy gene product crosses the threshold after 20 cycles of PCR.

**Supplementary Figure 4:** Dissociation curve obtained from MMQPCR showing two peaks - one for telomere product at 82°C and other at 86°C for single copy gene (albumin) product. A huge difference between the size of telomere and single copy gene peak can be seen.

**Supplementary Figure 5:** Standard curve generated for telomere product at step 2 (59°C) of stage 4 generated by MMQPCR with R^2^= 0.98, slope= -4.30 and efficiency of 70.7%.

**Supplementary Figure 6:** Standard curve generated for scg product at step 4 (85°C) of stage 4, generated by MMQPCR with R^2^= 0.99, slope= -3.08 and efficiency of 112.9%.

**Supplementary Figure 7:** Standard curve generated for single copy gene product at step (59°C) of stage 4, where the signal acquisition for telomere product had been captured, generated by MMQPCR with R^2^= 0.98, slope= -2.448 and efficiency of 156.2%.

**Supplementary Figure 8:** Representation of telomere amplification plots generated after performing 3-fold serial DNA dilution ranging from151ng-1.7ng DNA with MMQPCR methodology.

**Supplementary Figure 9:** Representation of single copy gene amplification plots at step 4 (85°C) of stage 4 generated after performing 3-fold serial DNA dilution ranging from151ng-1.7ng DNA with MMQPCR methodology.

**Supplementary Figure 10:** Representation of single copy gene amplification plots at step 2 (59°C) of stage 4 generated after performing 3-fold serial DNA dilution ranging from151ng-1.7ng DNA with MMQPCR methodology.

**Supplementary Figure 11:** 3% Agarose Gel was performed to analyze the product obtained from dual labeled fluorescence probe based assay; Telomere product was of 79bp and SCG (albumin) product was of 116bp. L1 represents 50bp ladder, L2 represents telomere product, L3 represents albumin product, L4 represents telomere product, L5 represents albumin product, L6 represents telomere product, L7 represents albumin product, L8 represents telomere product, L9 represents albumin product, L10 represents telomere product and L11 represents albumin product.

**Supplementary Figure 12:** Representation of amplification plots derived from dual labeled fluorescence probe based assay. A large difference of number of PCR cycles can be seen amongst telomere product and single copy gene product.

**Supplementary Figure 13:** Representation of amplification plots of telomere product, generated after performing 3-fold serial dilution with dual labeled fluorescence probe based assay methodology.

**Supplementary Figure 14:** Representation of amplification plots of single copy gene product, generated after performing 3-fold serial dilution with dual labeled fluorescence probe based assay methodology.

## REFERENCES

1. Sethi I. et al. Role of telomeres and associated maintenance genes in Type 2 Diabetes Mellitus: A review. Diabetes Research and Clinical Practice 122, 92–100.

2. Shalev I. et al. Stress and telomere biology: a lifespan perspective. Psychoneuroendocrinology 38, 1835–1842 (2013).

3. Eisenberg, D.T., Kuzawa, C.W. & Hayes, M.G. Improving qPCR telomere length assays: Controlling for well position effects increases statistical power. Am J Hum Biol 27, 570–575 (2015).

4. Aubert, G., Hills, M. & Lansdorp, P.M. Telomere length measurement-caveats and a critical assessment of the available technologies and tools. Mutation research 730, 59–67 (2012).

5. Montpetit A.J. et al. Telomere length: a review of methods for measurement. Nurs Res 63, 289-299 (2014).

6. Ding Z. et al. Estimating telomere length from whole genome sequence data. Nucleic Acids Research 42, e75–e75 (2014).

7. Cawthon R.M. Telomere length measurement by a novel monochrome multiplex quantitative PCR method. Nucleic acids research 37, e21 (2009).

8. Heid, C.A., Stevens, J., Livak, K.J. & Williams, P.M. Real time quantitative PCR. Genome research 6, 986–994 (1996).

9. Arya M. et al. Basic principles of real-time quantitative PCR. Expert Review of Molecular Diagnostics 5, 209–219 (2005).

10. Schmittgen T.D. & Livak K.J. Analyzing real-time PCR data by the comparative CT method. Nature Protocols 3, 1101 (2008).

11. Kibbe W.A. OligoCalc: an online oligonucleotide properties calculator. Nucleic acids research 35, W43–46 (2007).

12. Holland, P.M., Abramson, R.D., Watson, R. & Gelfand, D.H. Detection of specific polymerase chain reaction product by utilizing the 5’—3’ exonuclease activity of Thermus aquaticus DNA polymerase. Proceedings of the National Academy of Sciences of the United States of America 88, 7276–7280 (1991).

