## Supplementary Data_Sethi et.al for "Novel Dual Labeled Fluorescence Probe Based Assay to Measure the Telomere Length"

**† - Equal Authorship**

**Corresponding Author:**

Dr. Swarkar Sharma
Coordinator, Human Genetics Research Group,
School of Biotechnology, Shri Mata Vaishno Devi University, Katra, J&K, India


### ABSTRACT

Telomeres are highly repetitive regions capping the chromosomes and composed of multiple units of hexa-nucleotides, TTAGGG which makes their quantification difficult. Most of the methods developed to estimate telomeres are extensively cumbersome and expensive. The quantitative polymerase chain reaction (qPCR) based assay is relatively easy and cheaper method that applies SyBr Green dye chemistry to measure telomere length. As SyBr Green dye fluoresces after intercalation into the dsDNA, lack of differentiation between specific PCR target products and unspecific products is a limitation and it affects accuracy in quantitation of telomeres. To overcome the limitations of SyBr Green, we developed a dual labeled fluorescence probe based quantitative polymerase chain reaction (qPCR) to measure the telomere length. This robust, accurate and highly reproducible ($R^2$=0.96) proprietary method (patent pending), yet cost effective and easy, utilizes a probe that targets specifically the telomeric DNA.

### SUPPLEMENTARY TABLES

**Supplementary Table 1:** MMQPCR performed on 50 samples, the average delta Ct observed was -0.44. This implies the difference between copy number of telomere and single copy gene is very low.

| Samples | Number of Samples | Average ΔCt | Std. Dev. Ct |
|---|---|---|---|
| MMQPCR | 50 | -0.44 | 1.50 |

**Supplementary Table 2:** Comparison of both methodologies – MMQPCR and dual labeled fluorescence probe based assay for Telomere length measurement in a same sample set of 25. The samples were run in triplicates. Two of the samples did not provide the triplicate results hence removed from analysis. The copy number difference estimated by dual labeled fluorescence probe based assay between the telomere and scg was 5.04 folds as compared to 0.44 estimated by MMQPCR assay.

| Method | Number of Samples | Average ΔCt | Std. Dev. Ct |
|---|---|---|---|
| MMQPCR | 23 | -0.44 | 1.64 |
| Dual Labeled Fluorescence Probe based assay | 23 | -5.04 | 1.83 |

**Supplementary Table 3:** Telomere length Measurement by dual labeled fluorescence probe based assay in two independent cohorts of size n=120 and n=90 respectively, from population of Jammu and Kashmir, India. Samples were run in triplicates and averaged Ct values were used for each sample to estimate average ΔCt values ($\Delta Ct = Ct_{Telomere} - Ct_{Single\ copy\ gene}$).

| Method | Number of Samples | Average ΔCt | Std. Dev. Ct |
|---|---|---|---|
| Cohort 1 | 120 | -5.42 | 2.49 |
| Cohort 2 | 90 | -5.84 | 2.94 |

**SUPPLEMETARY FIGURES**

**Supplementary Figure 1:** 10% Native PAGE was performed to analyze the product obtained from MMQPCR; Telomere product was of 79bp and single copy gene product was of 98bp. L1 represents 50bp ladder, L2 represents albumin and telomere product, L3 represents telomere product, L4 represents albumin product, L5 represents albumin and telomere product, L6 represents telomere product, L7 represents albumin product and L8 represents NTC.

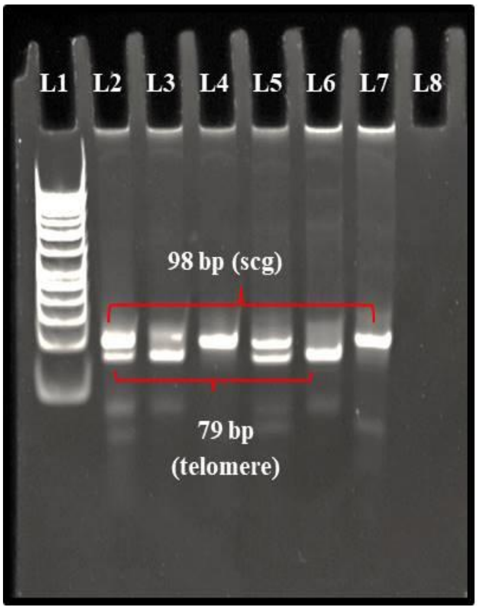


**Supplementary Figure 2:** Representation of amplification plot of telomere product obtained from MMQPCR. Telomere product crossed the threshold after 19 cycles of PCR.

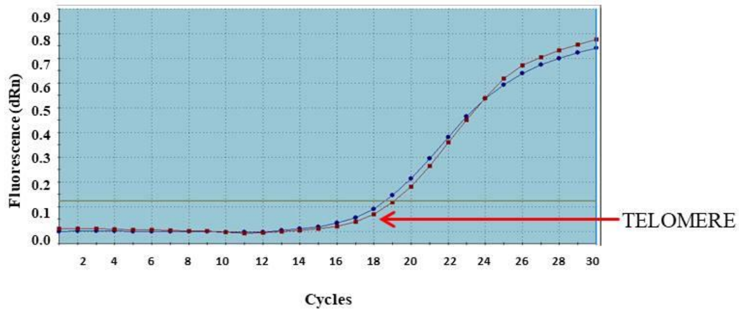

**Supplementary Figure 3:** Representation of amplification plot of Single Copy Gene (albumin) product obtained from MMQPCR. Single copy gene product crosses the threshold after 20 cycles of PCR.

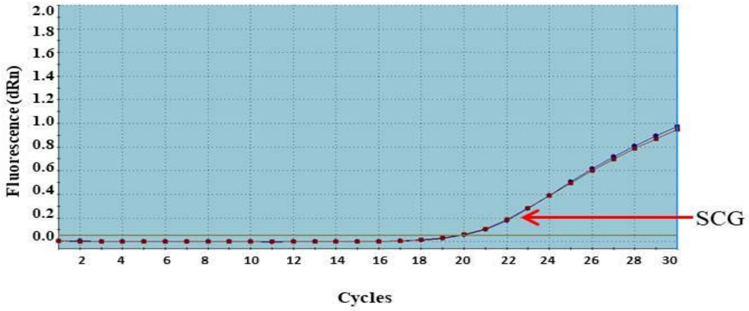


**Supplementary Figure 4:** Dissociation curve obtained from MMQPCR showing two peaks – one for telomere product at 82˚C and other at 86˚C for SCG (albumin) product. A huge difference between the size of telomere peak and single copy gene can be seen.

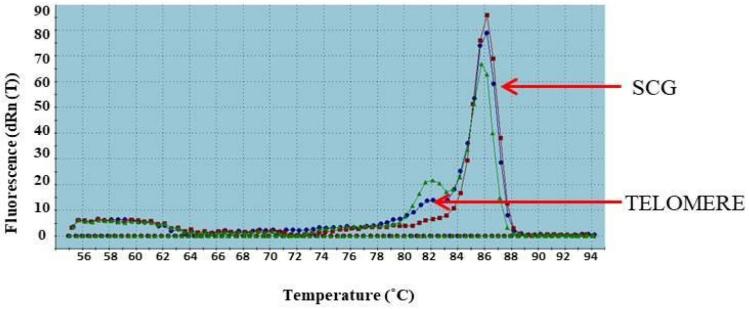


**Supplementary Figure 5:** Standard curve generated for telomere product at step 2 (59ºC) of stage 4 generated by MMQPCR with $R^2$= 0.98, slope= -4.30 and efficiency of 70.7%.

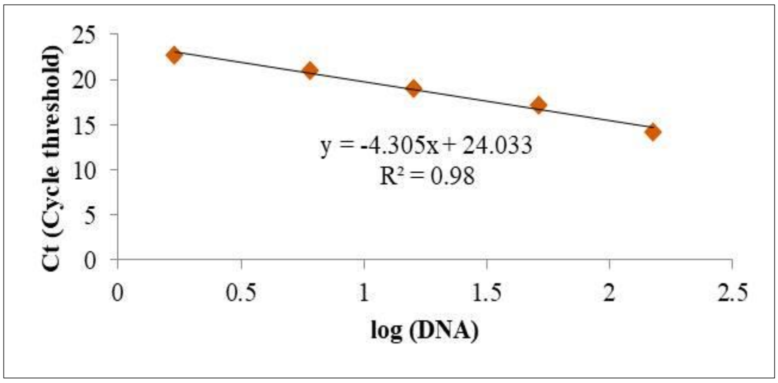

**Supplementary Figure 6:** Standard curve generated for scg product at step 4 (85ºC) of stage 4, generated by MMQPCR with $R^2$= 0.99, slope= -3.08 and efficiency of 112.9%.

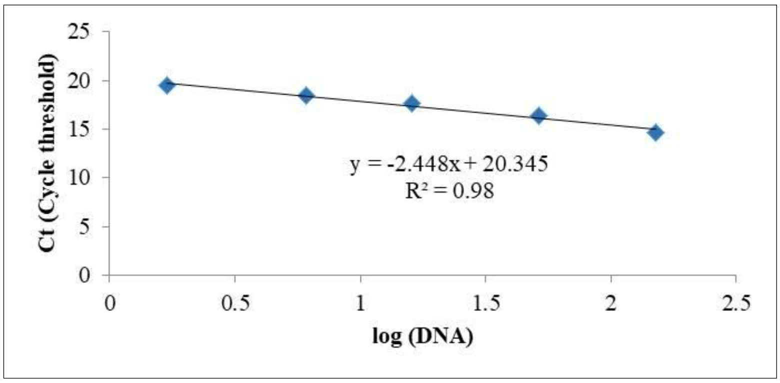


**Supplementary Figure 7:** Standard curve generated for scg product at step 2 (59ºC) of stage 4, where the signal acquisition for telomere product had been captured, generated by MMQPCR with $R^2$= 0.98, slope= -2.448 and efficiency of 156.2%.

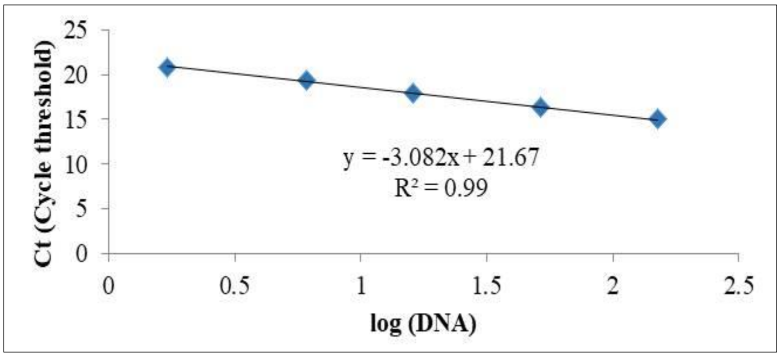


**Supplementary Figure 8:** Representation of telomere amplification plots generated after performing 3-fold serial DNA dilution ranging from151ng-1.7ng DNA with MMQPCR methodology.

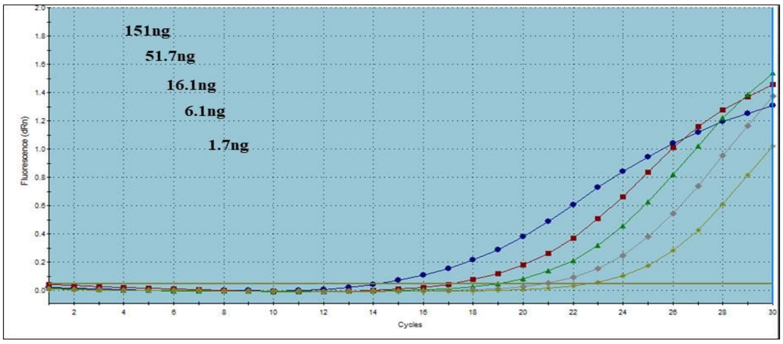

**Supplementary Figure 9:** Representation of scg amplification plots at step 4 (85ºC) of stage 4 generated after performing 3-fold serial DNA dilution ranging from151ng-1.7ng DNA with MMQPCR methodology.

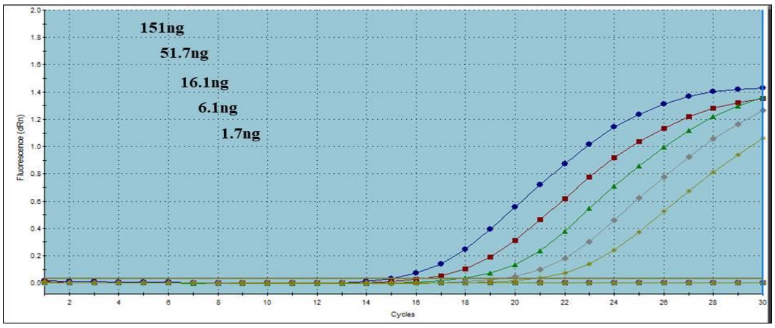


**Supplementary Figure 10:** Representation of scg amplification plots at step 2 (59ºC) of stage 4 generated after performing 3-fold serial DNA dilution ranging from151ng-1.7ng DNA with MMQPCR methodology.

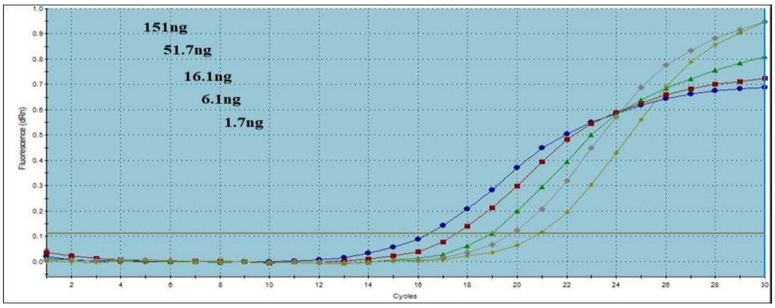

**Supplementary Figure 11:** 3% Agarose Gel was performed to analyze the product obtained from dual labeled fluorescence probe based assay; Telomere product was of 79bp and SCG (albumin) product was of 116bp. L1 represents 50bp ladder, L2 represents telomere product, L3 represents albumin product, L4 represents telomere product, L5 represents albumin product, L6 represents telomere product, L7 represents albumin product, L8 represents telomere product, L9 represents albumin product, L10 represents telomere product and L11 represents albumin product.

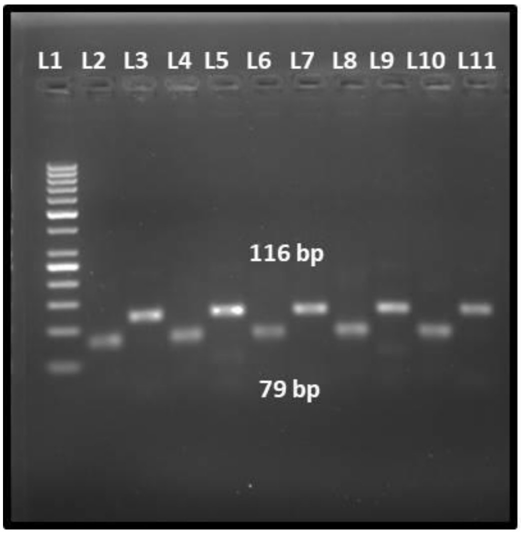


**Supplementary Figure 12:** Representation of amplification plots derived from dual labeled fluorescence probe based assay. A large difference of number of PCR cycles can be seen amongst telomere product and single copy gene product.

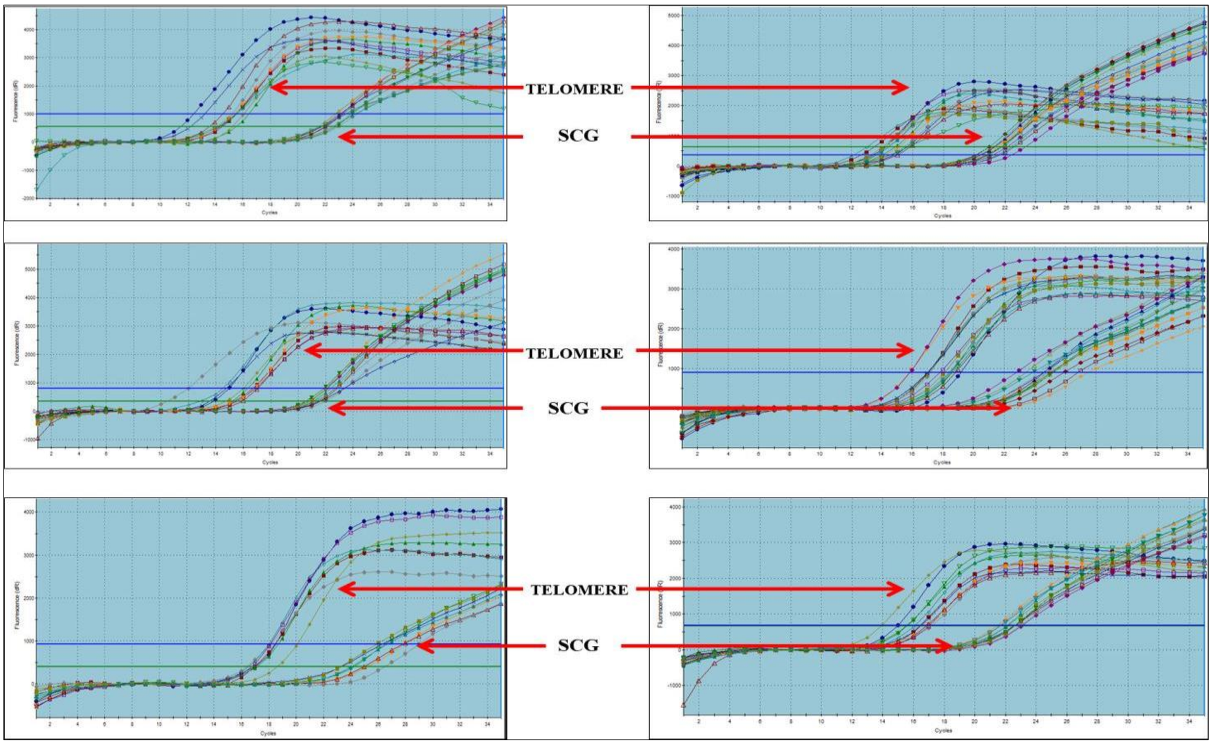

**Supplementary Figure 13:** Representation of amplification plots of telomere product, generated after performing 3-fold serial dilution with dual labeled fluorescence probe based assay methodology.

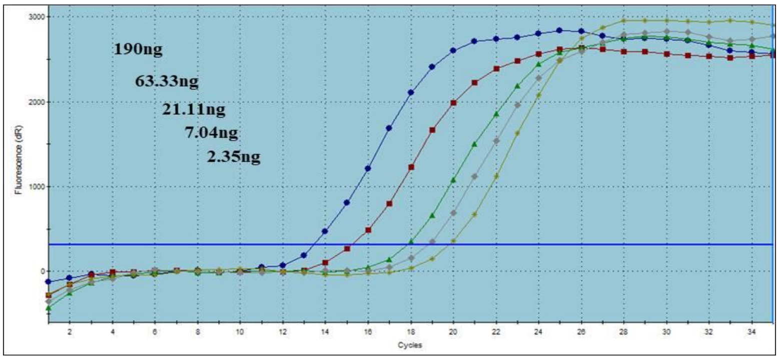


**Supplementary Figure 14:** Representation of amplification plots of scg product, generated after performing 3-fold serial dilution with dual labeled fluorescence probe based assay methodology.

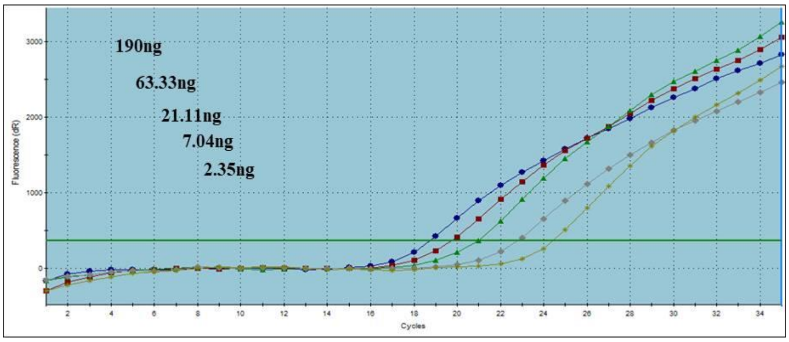
